# Real-time invasion dynamics reveal the drivers of predator spread and prey extirpation on an island

**DOI:** 10.64898/2025.12.22.695872

**Authors:** Guillem Casbas, Marc Vez-Garzón, Elba Montes, Victor Colomar, Oriol Lapiedra

## Abstract

Non-native predators profoundly restructure island biological communities worldwide, yet their management is hindered by limited empirical understanding of the dynamics of predator spread and prey extirpation. Here, we integrate two decades of field surveys, citizen-science records, and standardized transects to reconstruct the invasion of the snake *Hemorrhois hippocrepis* on Ibiza and its impact on the endangered, endemic lizard Podarcis pityusensis. We found a marked acceleration in range expansion despite intensified culling, evidence for the transition from establishment to spread stages of this invasion. Spatially explicit invasion maps combined with citizen reports of lizard disappearance show a non-linear acceleration in the interval from predator arrival to local prey extirpation, shortening from over ten years in early-invaded areas to three years in recently invaded areas. The observed patterns are consistent with density-dependent predator pressure, which generates a wave-like, rapidly advancing invasion front evidenced by declining snake body condition and reduced efficiency of snake captures in long-invaded areas. Our study empirically confirms theoretical predictions of nonlinear spread and impact dynamics and highlights how citizen science can crucially help documenting early invasion stages. Effectively incorporating the non-linear dynamics of invasions into conservation strategies is urgent to mitigate invasion-driven transformations of fragile island ecosystems worldwide.

## INTRODUCTION

Islands worldwide have become hotspots of biodiversity loss (Simberloff et al. 2013; Tershy et al. 2015; Bellard et al. 2016; Doherty et al. 2016; Spatz et al. 2017). Biological invasions, and invasive predators in particular, are the leading cause of this global phenomenon, having driven numerous local extirpations and global extinctions across island ecosystems worldwide (Salo et al. 2007; Sax et al. 2007; Medina et al. 2011; Doherty et al. 2016; Spatz et al. 2017). These biotic losses often result in irreversible functional and phylogenetic erosion in island biological communities (Kumschick et al. 2015; Matthews et al. 2024), with cascading effects on ecosystem services (Kamaru et al., 2024; Rogers et al., 2017; Tershy et al., 2015; Vila & Hulme, 2017). Managing invasive predators on islands is therefore an urgent priority for global biodiversity conservation in an era of rapid environmental changes (Doherty et al. 2016). Yet, effective management of invasive predators is largely limited by a lack of empirical data on key aspects of predator invasion dynamics (Epanchin-Niell and Hastings 2010; Baker and Bode 2016).

A major gap to effectively respond to the global threat of nonnative predators remains to disentangle the temporal dynamics of the spread and impact of predator invasions (Hulme 2009; Early et al. 2016). Because much of the damage to native communities occurs during the spread stage (Spatz et al. 2017), understanding the transition from establishment to spread is critical to tackle invasion progression (Blackburn et al. 2011). Reconstructing the fine-scale temporal dynamics of invasive predator spread requires analyzing long-term empirical datasets including extensive observations of the transition between the establishment and the spread stage of the invasion (Johnston and Purkis 2011). Consequently, studies uncovering both the dynamics and consequences of this transition in natural systems are scarce mostly due to the difficulty to track invasions in their early invasion stages as they unfold (Kolar and Lodge 2001). Indeed, data on the impact of alien predators often becomes available only once it is pervasive and after predators are widespread (Simberloff et al. 2013; Pyšek et al. 2020). Invasive predators are often not conspicuous and difficult to detect in the absence of ongoing local biodiversity tracking systems in the newly invaded area. This is especially true for snake invasions, with their commonly secretive behavior often obscuring patterns of establishment and spread (Christy et al. 2010). Consequently, snake invasions often wreak havoc in native biological communities worldwide (Savidge 1987; Dorcas et al. 2012; Kraus 2015; Hinckley et al. 2017; Pyšek et al. 2020; Montes et al. 2021; Piquet and López-Darias 2021).

A complementary unresolved, critical question in invasion biology is to untangle the dynamics of native species decline and extirpation following the spread of invasive predators. Reliable data on this process is however scarce since it requires high-quality data on prey dynamics before, during, and after the arrival of invasive predators. Only by explicitly linking the range expansion of invasive predators to robust long-term data on the decline, range contraction, or local extirpation of their native prey can we fully understand the dynamics of this process (Meyerson et al. 2019). However, empirical studies coupling these two processes remain scarce and focused on post-impact scenarios (as highlighted by Doherty et al., (2016)). Untangling the real-time impact dynamics of invasive predators on native prey has therefore become a global conservation priority to guide targeted conservation actions to minimize biodiversity loss (Jeschke and Strayer 2008; Piquet and López-Darias 2021) and preserve ecosystem functioning, particularly on islands (Doherty et al., 2016; Kraus, 2015; Rogers et al. 2017).

To address this question, we leveraged a rare opportunity to study predator spread and prey extirpation in real time. In the Mediterranean island of Ibiza, a rapidly spreading invasive horseshoe whip snake is extirpating an endangered endemic Ibiza wall lizard (Hinckley et al. 2017; Montes et al. 2021). By integrating two decades of data from thousands of geo-referenced snake captures and lizard observations on Ibiza with standardized field transects and different sources of citizen science data, here we first characterize the spatial and temporal progression of the invasive snake’s range, with a particular focus in untangling the transition between establishment and spread stages. We then examine if there is a tight temporal association between predator arrival and local extirpation of the endemic lizard. Then, we assess how these dynamics change over time by characterizing the acceleration of extirpation as a stronger invasion front develops. Finally, we examine whether and how food depletion is linked to lower predator abundance at the core of the invasion as well as to the emergence of an invasion front. Our findings provide new insights into both the patterns and processes of invasive predator spread dynamics as well as the rapid extirpation of endemic species that have wide implications for conservation and management of worldwide island biodiversity under global change.

## METHODS

### 1. Study system and invasion history

The study was conducted on the island of Ibiza (572.6 km²), part of the Balearic archipelago in the western Mediterranean. Ibiza is characterized by a mosaic of dry shrublands, pine forests, and agricultural land, prime habitat for both the invasive predator and its endemic prey (Salvador 2009; Feriche 2017). The invasion of the horseshoe whip snake was first confirmed in 2003, when an individual emerged from the hollow trunk of an ornamental olive tree imported from mainland Spain (Servei d’Agents de Medi Ambient 2003). From this very localized distribution, the invasion rapidly spread across most of the island in a 20-year period (Dappen et al. 2013; Montes et al. 2021). By 2023, snakes were present in 83,36% of the island (Fig. 1) despite management efforts including intensive trapping campaigns.

**Figure 1.**
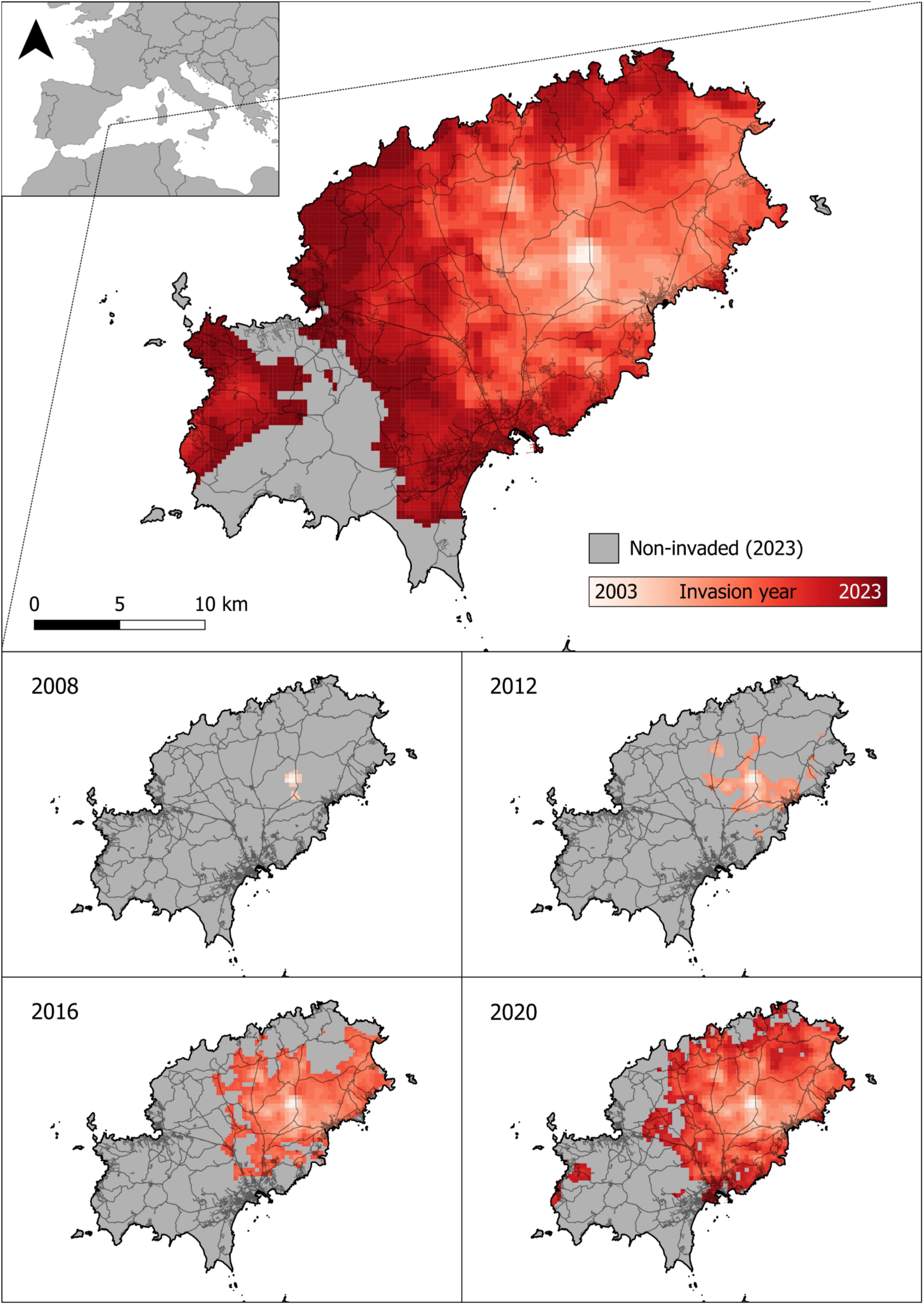
Chronology of snake invasion across Ibiza as reconstructed using empirical and citizen science data. The panel shows the estimated year of the horseshoe whip snake presence across the island. Warmer colors indicate earlier invasion years. Grey cells represent areas without confirmed invasion as of 2023 (top). Bottom panels show detailed invasion fronts at four key time points (2008, 2012, 2016, and 2020).

The horseshoe whip snake is a diurnal, non-venomous colubrid native to North Africa and the Iberian Peninsula. It is a ground-dwelling, active predator that thrives on Mediterranean shrublands, with lizards representing an important part of their diet in their native range (Pleguezuelos and Moreno 1990). Culling of snakes in Ibiza began early in the invasion, yielding almost 6,000 capture records with associated geographic information (Supplementary Fig. S1) across two decades.

The endemic Ibiza wall lizard is an icon of local culture and plays a keystone role in the island biological community as a primary insect consumer, pollinator and seed disperser (Traveset 1995; Hinckley et al. 2017). This lizard composes over 55% of prey items of the horseshoe whip snake in Ibiza (Hinckley et al. 2017). Given their rapid decline linked to intense predation by invasive snakes, it was recently re-classified as Endangered by the IUCN in 2024 (Bowles 2024). The island tameness of this species makes it particularly vulnerable to snake-driven extirpation, as Ibiza had no native snake species prior to this introduction (Silva-Rocha et al. 2015). Being a tame, beloved, and culturally iconic species (Pérez Mellado 2009; Dappen et al. 2013) we were able to reconstruct its disappearance using local knowledge collected though structured citizen science surveys (Mohanty and Measey 2019) complemented with standardized field transects.

### 2. Mapping invasion spread through time

To reconstruct the spatiotemporal dynamics of the horseshoe whip snake invasion in Ibiza, we compiled two types of data spanning two decades (see table S1): (i) empirical snake occurrence records and (ii) citizen science observations. Empirical snake occurrence data points were obtained from long-term trapping efforts from an eradication program (n = 4394), roadkill detections (n = 77), and integrated field records reported in Montes et al. 2021 (n = 1271). We complemented this dataset by using citizen science data surveys. Thanks to the human innate predisposition to respond strongly to snakes (Öhman and Mineka 2001) and the herpeto-phobic attitude of many Mediterranean cultures (Silva-Rocha et al. 2015), residents often clearly recall their first encounter with a snake. These reports provided valuable temporal and spatial coverage for reconstructing the early stages of the invasion. They included targeted in-person surveys with residents of rural households (n = 60), responses to an online snake sighting questionnaire (n = 172), and validated observations submitted to iNaturalist (n = 14).

In parallel, we also collected 120 citizen reports from participants who stated they had never observed snakes in their area. While such reported absences may reflect either true absence or a lack of detection, they provided a useful reference to delineate the limits of the invasion front at different time points. We treated these data conservatively, considering them provisional absences unless contradicted by alternative sources considered to be more reliable such as actual snake captures. When direct observations—such as roadkills or captures—occurred in the same location and year, they were used to cross-validate and, when necessary, override both reported sightings and absences. Reports with conflicting dates or vague spatial information were excluded.

Finally, to detect early snake presence in areas just ahead of the invasion front, we installed dozens of ‘sentinel’ traps at sites where snakes had not yet been confirmed. These traps served as a proactive tool to monitor the leading edge of the invasion and provided real-time information on geographic expansion.

To generate the map of snake spatial distribution through time, we used QGIS 3.28 to process all spatial data. We first buffered each georeferenced snake occurrence point with a 500 m radius. Then, we overlaid this information on a 500 × 500 m grid covering the whole island, where each cell was assigned the earliest year of confirmed snake presence. This resulted in a dynamic temporal map that included information for 79.8% of the island 500x500m grid cells (this is, n = 1987 grid cells with invasion data out of a total of 2493 grid cells). To estimate year of invasion in unsampled areas and to smooth abrupt transitions between detection points, we applied inverse distance weighting (IDW) interpolation (p = 5). We used the resulting dynamic raster file to estimate the spatiotemporal progression of horseshoe whip snake throughout the island from 2003 to 2023.

### 3. Quantifying expansion dynamics

We used the dynamic raster file to calculate the cumulative area invaded annually. To do so, we extracted the total area classified as invaded each year directly from the raster map, as shown in Fig. 2C. To quantify the invasion speed, we traced 16 cardinal axes from the introduction point (Supplementary Fig. S2). Then, we recorded distances reached by the invasion front year-by-year along each of these axis, and excluded from each yearly estimate of expansion those axes that had reached the coastline the previous year and therefore could not expand further. We averaged new distance covered by the invasion across all remaining axes, after excluding those axes that had reached the coast. Similarly, we calculated invasion progress speed (distance/year) to assess whether spread rates were constant or accelerating. Finally, we used a segmented regression analysis, implemented via the ‘*segmented’* R package (Muggeo 2008) to characterize changes in the slope of invasion expansion over time to detect statistically significant slowdowns or accelerations such as the transition between establishment and spread phases (Fig 2A).

**Figure 2.**
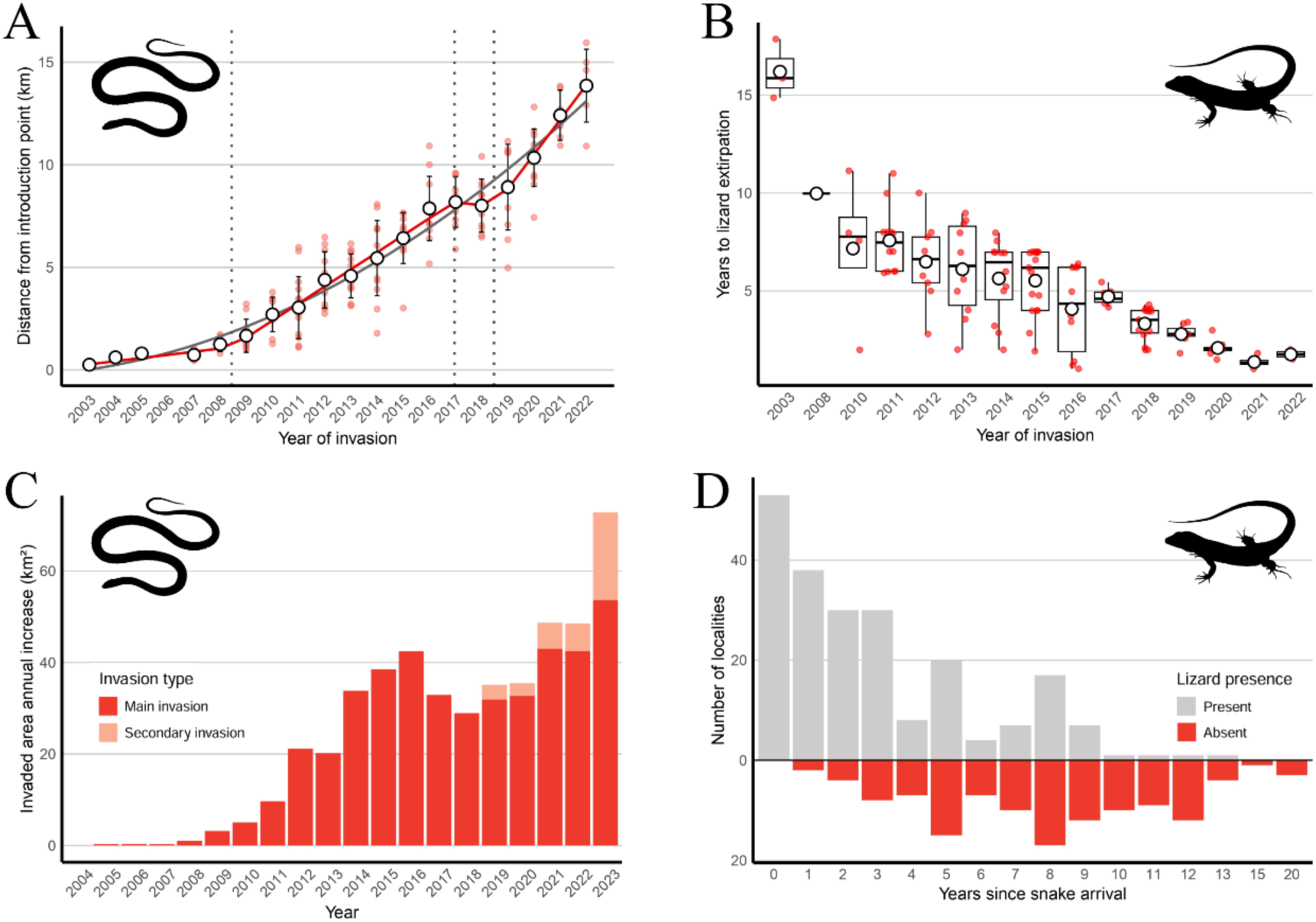
Spatiotemporal dynamics of the snake invasion and associated lizard extirpation patterns in Ibiza. (A) Distance from the introduction point over time along 16 cardinal directions, showing nonlinear acceleration in range expansion. Dotted vertical lines indicate breakpoints separating invasion stages. The solid red line indicates the mean slope of each invasion phase, and the solid grey line, the polynomic adjustment. (B) Relationship between year of snake arrival and time to local lizard extirpation. (C) Cumulative area invaded by snakes (km²) from 2003 to 2023, differentiating between main invasion (dark) and secondary invasion (light) (D) Histogram of time since snake arrival in all surveyed grid cells, differentiating sites where lizards are still present (grey) versus extirpated (red).

### 4. Timing local lizard disappearance across the invasion

To examine the impact of snake invasion on the Ibiza wall lizard population, we calculated the time lag between the first estimated snake presence, and the last year lizards were observed at each location. To estimate local extirpation dates we used two data sources: (i) structured citizen science surveys targeted to rural households across the island, from which we collected a total of 358 responses; of these, 219 were excluded because lizards were still reported as present, and 35 were excluded due to time inconsistencies or due to missing data, resulting in 104 valid records; and (ii) complementary field surveys to actively search for lizards in locations that were frequently visited for other research projects or to monitor lizard presence (n = 17 of sites surveyed repeatedly across several years until lizards were no longer detected). Then, we tested whether the time between snake arrival and lizard disappearance became shorter in more recently invaded areas by plotting extirpation time lags against the year of invasion and fitting a linear regression model (Fig. 2B).

### 5. Assessing prey depletion along the invasion

We used three complementary approaches to evaluate whether and how prey depletion shapes the dynamics of the invasion and contributes to create a wave-like pattern at the invasion front. First, we tested whether snakes in long-invaded areas are in worse body condition than those at the forefront of the invasion. To test this hypothesis, we collected 1505 snakes between 2009 and 2023 from the ongoing culling campaign in Ibiza and measured their SVL and body mass. We overlapped the geo-referenced location of each specific capture to the year of invasion at that specific location. Then, we built a linear model to assess if the year of invasion was associated to body condition measured as the division of log (body weight) by the log (snout–vent length) (Fig. 3B, Table S2).

**Figure 3.**
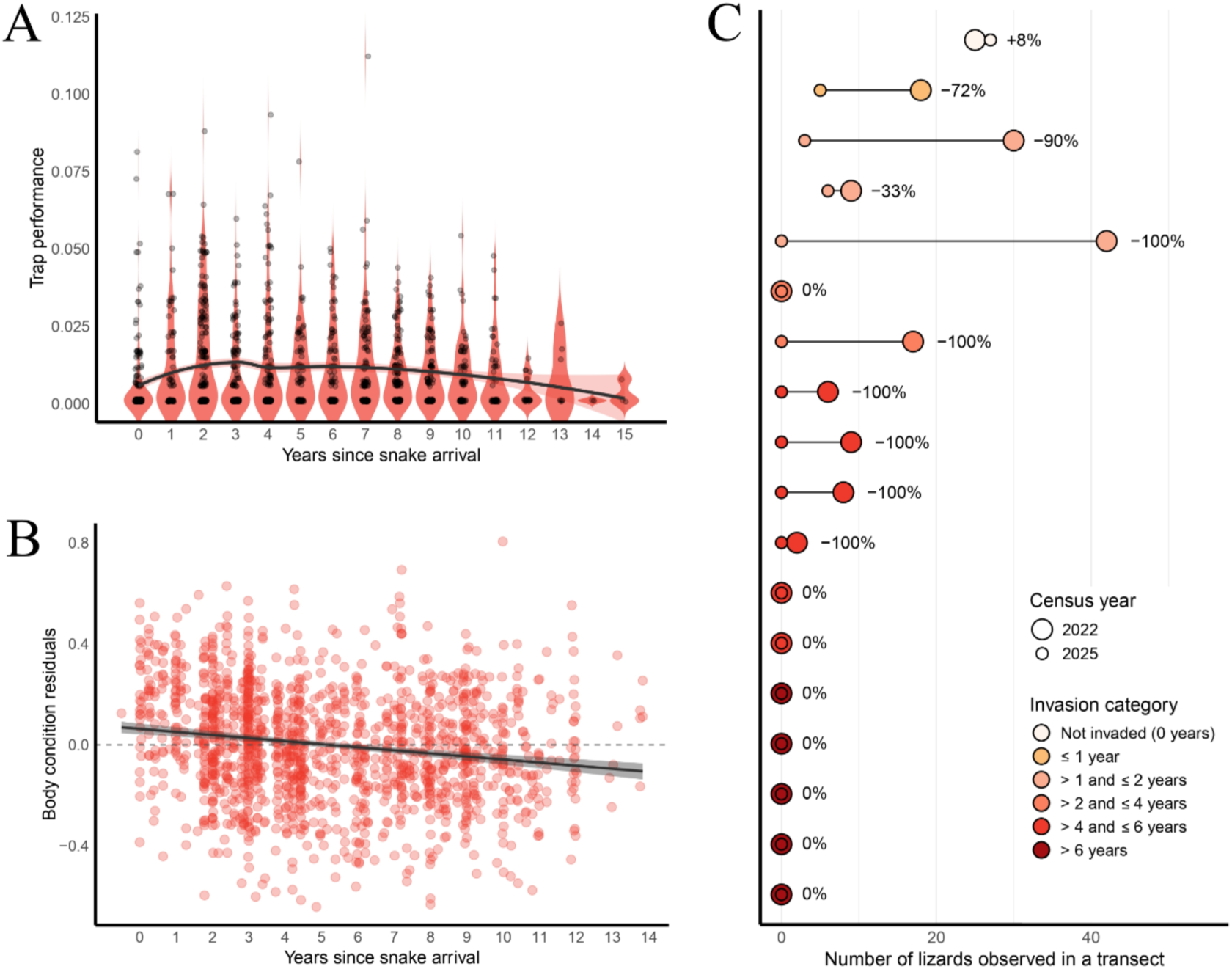
Evidence for prey depletion and associated effects of prolonged snake invasion. (A) Trap performance (captures per trap-night) plotted against the number of years since snake arrival. (B) Body condition of snakes, represented as the residuals of the log(mass) ∼ log(SVL), plotted against years since invasion. (C) Lizard abundance per transect from 2022 to 2025 across invasion categories. Circle size reflects the number of lizards recorded; color indicates the number of years since snake arrival at each site (as of 2025). Lines connect data points from the same site across years.

Second, we assessed changes in lizard abundance using standardized visual encounter surveys conducted at 18 sites both in 2022 and 2025. At each site, we surveyed between two and four fixed transects each year, with an average length of 332 meters (±111 m SD), following the same routes across years. Transects followed traditional drystone walls in semi-abandoned Mediterranean agricultural landscapes. During each survey, lizards were recorded only when observed within a 3-meter band directly facing the wall. This habitat and method were selected to minimize site-to-site variability and standardize detection conditions. For each site and year, we used the maximum number of lizards recorded across repeated visits as a relative abundance estimate. In total, the visual encounter surveys included 77 transects across two years and 18 sites, covering 17,319 m² and resulting in 207 lizard observations. In 2022, the 18 study sites spanned different invasion stages: 7 sites were already invaded and had no lizards, 5 sites had lizards but no snake presence, and 6 sites had both snakes and lizards. By 2025, all sites had been invaded by snakes, and only 4 still harbored lizard populations. This progression allowed us to track changes in lizard abundance over time within the same transects as the invasion advanced.

Third, we quantified trap capture efficiency as the number of snakes captured per trap-day, excluding the winter season (November to April) when traps were inactive. We analyzed data from 4,981 geo-referenced traps deployed across Ibiza between 2016 and 2023, totaling over 4.17 million trap-days (4,176,631 days in total), calculated as the number of days each trap was active during that period. This large-scale dataset reflects a substantial management effort and offers a powerful view into predator dynamics over time. Then, we evaluated whether snake capture efficiency decreased as a function of years since invasion by modeling trap efficiency data using a zero-inflated mixed-effects model implemented in the ‘*glmmTMB’* package (Brooks et al. 2017). Model assumptions were assessed using the ‘*performance’* package (Lüdecke et al. 2021). This model allowed us to estimate both the probability of zero captures and the average trap efficiency across invasion time (Fig.3A, Table S3).

## RESULTS

### Predator range acceleration defines the transition between establishment and spread

Our map of snake invasion shows that up to 2023 snakes had spread to almost the totality of the area of the island (83.4 %, Fig 1C). Through time, the progress of the invasion front across 16 cardinal points shows a non-linear spread. Specifically, the pattern of snake expansion through time best fits a polynomic function adjustment (F-statistic = 756.5, df = 193, p < 0.001, R2 = 0.8857) (Fig 1; Fig 2A).

We detected three breaking points where the slope of average geographical expansion rate significantly shifts (Fig 2A), defining four different stages of invasion. A first phase of very limited range expansion during the first 5 years after detected arrival of snakes in 2003 could be described as the establishment phase of the invasion where, in fact, the slope does not differ from zero (estimate = 0.154 km/year, std. error = 0.093, t value = 1.654, CI = [-0.029, 0.338]. Then, we detected a sharp increase in expansion rates in 2008.4 (+/- 0.6 years). This significant change in the slope (estimate = 0.831 km/year, std. error = 0.057, t value = 14.511, CI = [0.718, 0.944] (Fig 2A) defines the transition between the establishment and spread phases of this predator invasion. Unexpectedly, we found a significant slowdown in expansion rate in 2016.9 (+/- 0.5 years) (estimate = -0.175 km/year, std. error = 0.561, t value = -0.312, CI = [-1.282, 0.931]). This is coincident with the invasion reaching a major road with heavy traffic in its western (south-west to north-west) direction (Ministerio de Transportes 2019). The expansion rate, however, accelerated again in 2018.5 (+/- 0.4 years, (estimate = 1.687 km/year, std. error = 0.199, t value = 8.461, CI = [1.294, 2.081] Fig 2B). This point established the beginning of a new rapid expansion phase that is still ongoing. Interestingly, a recent reinforcement of culling effort transitioning from 223 traps in 2018 to over 1.200 after 2021 did not result in a new decrease in the rate of geographical expansion.

### Local extirpation of the endemic Ibiza wall lizard estimated with citizen science data

By overlapping maps of invasion spread with those of lizard local extirpation our results show both processes to be tightly associated. Specifically, all lizard extirpations reported follow the establishment of snakes in each specific area. On average, extirpation occurred 5.18 ± 2.99 SD years after snake arrival (Fig. 2D). Time to local extirpation of the Ibiza wall lizard significantly decreased with later invasion years (β = –0.66 ± 0.04 SE, t = – 15.14, p < 0.001; R² = 0.66; AIC = 480.5), indicating that lizards disappeared more rapidly in recently invaded areas (Fig. 2B). A log-transformed model had a p < 0.001 and yielded a lower AIC (130), supporting an exponential acceleration in extirpation timing across the invasion timeline.

### Enhanced invasion expansion drives accelerated extirpation of its main prey

Patterns of local extirpation accelerated with time from the beginning of the invasion. In early phase of the invasion (locations invaded between 2003 and 2011), local lizard extirpation took on average 8.9 ± 3,78 SD years following snake arrival (Fig. 2B). This temporal pattern mostly corresponds to the establishment phase and the beginning of the spread phase of the snake invasion. In more recent years, during the next stage of the invasion, extirpation occurred faster. For sites invaded between 2012 and 2015, for instance, average time to extirpation was of 5.03 ± 1.87 SD years. Finally, extirpation happened as fast as within 3.19 ± 1.45 SD years in the most recently invaded sites (for instance in the case of sites invaded between 2016 and 2019; Fig. 2B). This accelerated impact to native prey populations is particularly striking given the intensive and increasing culling effort to eliminate snakes, particularly in recent years, where as many as ∼6000 snakes are estimated to be culled each year in this 572.6 km2 island.

### Prey depletion is associated with the emergence of an invasion front

We found evidence that food depletion is associated with the emergence of an invasion front where invasive predators are more abundant at the front of invasion than in its core. Snakes remaining in areas invaded for years are in significantly worse body condition than those in more recently invaded areas (β = −0.0449 ± 0.0062 SE, t = −7.23, p = 7.7 × 10⁻¹³, R² = 0.033, n = 1,513; Fig. 3B). Additionally, standardized visual surveys conducted in 18 sites across the invasion gradient revealed a sharp decline in relative lizard abundance in recently invaded sites, with an average reduction of 84,05% in observed individuals between 2022 and 2025 on those sites than still had lizards present in 2022 (Fig 3C). Parallel to a worsened body condition, our thousands of traps installed throughout the island revealed that predator capture efficiency significantly declined in areas that had been invaded for a longer time (Fig 3A, Table S3) (ZI: odds ratio = -1.58, CI = [-2.32, -0.84], *χ*^2^1, 0.05 = 17.60, p < 0.001; Cond.: odds ratio = -0.31, CI = [-0.54, -- 0.07], *χ*^2^1, 0.05 = 6.56, p = 0.01). This finding demonstrates lower snake densities over time as the amount of food available to snakes rapidly decreases in these long-invaded areas, generating the classical wave-like predicted pattern of expanding biological invasions (Fig 3A).

## DISCUSSION

Overwhelming evidence shows that predator invasions are global drivers of island biodiversity loss (e.g. (Savidge 1987; Salo et al. 2007; Medina et al. 2011; Tershy et al. 2015; Bellard et al. 2016; Russell et al. 2016; Doherty et al. 2016; Spatz et al. 2017; Vila and Hulme 2017; Rogers et al. 2017; Kamaru et al. 2024). However, our lack of understanding of key aspects of their invasion dynamics is a major stumbling block hindering their effective management worldwide (Hulme 2009; Epanchin-Niell and Hastings 2010; Baker and Bode 2016; Russell et al. 2016; Early et al. 2016). Indeed, a major challenge to disentangle the dynamics of invasive-predator-native-prey interactions as they unfold has been to compile detailed real-time, replicated empirical data of this process (Blackburn 2019). Our study empirically links the spatiotemporal dynamics of invasive predator spread to the collapse of an endemic prey species on an island, offering new insights into the processes driving biodiversity loss in fragile insular ecosystems worldwide.

By reconstructing the 20-year progression of the horseshoe whip snake invasion in Ibiza we contribute filling persistent gaps to unravel the dynamics of biological invasions. Specifically, we show how predator spread has led to the rapid collapse of an endemic keystone species and associate the emergence of an invasion front to a rapid resource depletion driven by the invasive predator itself. These findings contribute to identify the drivers of the transition from invasion establishment to spread, as well to accurately characterize the temporal dynamics of native species collapse following predator arrival.

### Unraveling the dynamics of invasive predator spread

Our study represents a high-resolution reconstruction of the invasion dynamics of the horseshoe whip snake on the Mediterranean island of Ibiza. By integrating long-term culling records, citizen science observations, and standardized field transects we identified a critical transition from the establishment phase to a rapid spread phase, marked by a sharp acceleration in geographic expansion despite intensified culling efforts. This transition aligns with theoretical models that predict nonlinear invasion dynamics and underscores the importance of early detection and intervention in managing invasive species (Epanchin-Niell and Hastings 2010; Blackburn et al. 2011). In addition, we hypothesized that covid-19 lockdown may have contributed to the acceleration phase after 2020. However, we found this acceleration was already detected before 2020 in our study area. Prior to this, we detected a deceleration of the invasion around 2017 which can likely be attributable to the invasion front reaching the main traffic artery of the island (Fig 1, Fig 2A).

Identifying the transition between establishment and spread can be crucial for informing management strategies, as it represents a window of opportunity where containment and eradication efforts are most likely to succeed. Our findings emphasize the need for continuous monitoring and adaptive management approaches that can respond swiftly to changes in invasion dynamics, particularly through early action before the spread phase begins.

### Linking predator spread to native species collapse

A central result of our study is the characterization of the temporal lag between predator establishment and prey extirpation, which consistently shortens over time. This indicates that the impact of the invasive predator intensifies as the invasion advances. The delay between snake establishment and lizard disappearance declined from around a decade in early-invaded sites to less than three years in recent ones in a period of just over 10 years (Fig 2B). These findings support theoretical predictions of non-linear and density-dependent invasion impacts (Taylor and Hastings 2005; Simberloff et al. 2013), but empirical confirmation of these dynamics has remained limited. Our results show how these processes play out in real time, highlighting the urgent need to intervene during the early stages of predator establishment before accelerated collapse becomes inevitable.

The spread dynamics of an invasion are not temporally or spatially static phenomena. Rather, they are emergent outcomes of how populations respond to the ecological conditions they experience (Meyerson et al. 2019). The accelerated invasion dynamics observed in our study system can result from different, non-mutually exclusive ecological processes. One possibility is that growing predator densities intensify per capita pressure on native prey populations, rapidly driving them below viability thresholds. Another factor is the ecological naivety of island prey to novel predators, which results in ineffective antipredator responses and little behavioral adjustment over short timescales. In addition, predators might be rapidly plastically or evolutionary adapting to new conditions in their invasive range, thereby becoming more efficient predators of native fauna. Combined, these factors highlight the vulnerability of island biotas to predation-driven collapse if invasive predators are not rapidly and effectively managed (Jeschke and Strayer 2008; Doherty et al. 2016)

### The global threat of invasive snakes

Invasive snakes have emerged as major drivers of biodiversity loss on islands, where they have become extremely efficient invasive predators (Savidge 1987; Wiles et al. 2003; Dorcas et al. 2012; Piquet and López-Darias 2021). This can be in part explained by the fact that snakes can survive for extended periods of time with little food income (McCue 2007). This buffers small population size stochastic events such as Allee effects (Kramer et al. 2017), which are an important cause of local extinction of nonnative species arriving to novel areas (Taylor and Hastings 2005; Kanarek and Webb 2010). Our findings from Ibiza resonate with other documented cases worldwide. For instance, the introduction of the brown tree snake (*Boiga irregularis*) to Guam led to the extirpation of most native forest bird species, resulting in cascading ecological effects, including disrupted seed dispersal and altered forest composition (Rogers et al. 2017). Similarly, the Burmese python (*Python bivittatus*) in Florida’s Everglades has caused severe declines in mammal populations, with some species experiencing reductions of over 90% (Dorcas et al. 2012). In the Canary Islands, the California kingsnake (*Lampropeltis californiae*) has decimated endemic reptile populations on Gran Canaria, with reductions exceeding 90% for some species, leading to significant ecological imbalances (Piquet and López-Darias 2021). These examples underscore the profound and often irreversible impacts invasive snakes can have on island ecosystems. Our study adds to this body of evidence by demonstrating the rapid and accelerating effects of snake invasions on native prey species, emphasizing the need for vigilant biosecurity measures and rapid response strategies to prevent similar outcomes in other vulnerable regions.

### Invasion fronts and their resource-driven expansion

We also provide evidence that prey depletion behind the invasion front may contribute to both the structure and accelerated spread of the invasion. Declines in trapping efficiency and snake body condition in long-invaded areas suggest that food resources, including the endemic lizard, become exhausted as the invasion proceeds. This supports the idea that invasive snakes are not only expanding opportunistically but are being ecologically pushed forward by the degradation of foraging conditions in already-colonized habitats. Such resource-driven invasion fronts represent a self-reinforcing mechanism of spatial spread and ecological disruption, in line with the predictions of consumer–resource models (Sherratt 2001).

These findings have clear management implications. Control measures focusing solely on suppressing predator abundance in long-invaded areas may not be sufficient to limit predator spatial spread or conserve threatened prey. Instead, adaptive strategies that anticipate and intercept invasion fronts, ideally during the transition from establishment to spread, may be more effective in minimizing impacts. This is particularly relevant given that the final wave of expansion in Ibiza occurred despite a tenfold increase in trapping effort, highlighting the difficulty of halting invasion momentum once underway.

### The value of citizen science in tracking invasions

Our study also demonstrates the critical role that citizen science can play in reconstructing invasion histories and documenting biodiversity impacts in real time. In many regions, especially on islands, professional monitoring networks are sparse or absent. In such contexts, data provided by local residents, landowners, and nature enthusiasts can offer a valuable complement to formal scientific monitoring (Mohanty and Measey 2019). These data can be especially useful in the case of iconic species such as the Ibiza wall lizard (Dappen et al., 2013). Indeed, beyond their informational value, these observations also reflect strong local concern and cultural attachment to this iconic declining species, which can be effectively used to provide empirical data critical to support conservation interventions. Citizen observations of lizard disappearances aligned closely with our field data, validating the reliability of such participatory approaches. This underscores the potential of citizen science as a cost-effective and scalable tool for monitoring invasive species and their impacts, particularly in regions where resources for extensive fieldwork may be limited (Crall et al. 2013).

### The importance of untangling invasion dynamics to protect native biodiversity in a rapidly changing planet

The dynamics we document in Ibiza illustrate how rapidly invasive species can restructure ecological communities, especially on islands where native species often lack the tools to effectively respond to them (Blumstein and Daniel 2005). More broadly, our findings emphasize the importance of integrating dynamic invasion processes, including feedbacks between resource availability, predator capture performance, and expansion dynamics, into biodiversity conservation planning. Indeed, our results contribute to a growing recognition that understanding the tempo of biological invasions is as critical as understanding their spatial extent. This is particularly urgent in a rapidly changing world, where global trade continues to increase the frequency and ecological footprint of biological invasions (Helmus et al. 2014; Bellard et al. 2016; Early et al. 2016; Seebens et al. 2017).

By elucidating the transition from establishment to rapid spread and documenting the acceleration of prey extirpation, we highlight critical phases where management interventions on predator invaders worldwide can be most effective. The acceleration of both range expansion and ecological impact in our study system suggests that windows for effective intervention may be narrower than previously assumed. Therefore, management strategies that prioritize early detection and rapid intervention are essential to confront the growing threat of invasive predators (Epanchin-Niell and Hastings 2010; Early et al. 2016; Meyerson et al. 2019). These insights are particularly relevant for island ecosystems, which are hotspots of biodiversity disproportionately affected by the negative impact of invasive species (Bellard et al., 2016; Spatz et al., 2017). Our findings underscore the necessity of integrating invasion dynamics into conservation planning and policy-making to mitigate the impacts of invasive predators and preserve ecosystem services in the current scenario of rapid, global environmental change.

## Acknowledgments

We thank GEN-GOB Eivissa for their invaluable support in conducting surveys and assisting with data collection from local residents, as well as for their continuous help with field logistics and overall support during our time on the island. We are also grateful to Joan Costa Vila for his generous assistance during fieldwork and for helping distribute the survey and engage with the local community.

## Author contributions (CRediT)

Guillem Casbas, Marc Vez-Garzón, and Oriol Lapiedra contributed to the conceptualization of the study, data curation, formal analysis, methodology design, visualization, and original draft preparation and writing. Guillem Casbas and Marc Vez-Garzón also contributed to funding acquisition and resource collection. Investigation was carried out by Guillem Casbas, Marc Vez-Garzón, Oriol Lapiedra, Víctor Colomar, and Elba Montes. Project administration, funding and supervision were led by Oriol Lapiedra. Resources were provided by Víctor Colomar, Elba Montes, Guillem Casbas, and Marc Vez-Garzón. Validation was performed by Oriol Lapiedra, Elba Montes, and Víctor Colomar. Writing – review and editing was carried out by Oriol Lapiedra, Elba Montes, Marc Vez-Garzón, Guillem Casbas and Víctor Colomar.

