## Supplementary material for "Real-time invasion dynamics reveal the drivers of predator spread and prey extirpation on an island"

| ***Data type*** | ***Source*** | ***Number of records*** | ***Years*** |
|---|---|---|---|
| *Snake occurrences* | COFIB eradication program | 4395 | 2016–2023 |
| *Snake occurrences* | Montes et al. 2021 | 1271 | 2010–2018 |
| *Snake occurrences* | Roadkill detections | 77 | 2021–2022 |
| *Citizen science* | In person rural household survey | 60 | 2008–2023 |
| *Citizen science* | Online survey (retained responses) | 172 | 2008–2023 |
| *Citizen science* | iNaturalist records | 14 | 2016–2023 |

**Table S1.** Data sources used to reconstruct the temporal invasion of the horseshoe whip snake in Ibiza.

| Term | Estimate | Std. Error | t value | p value |
|---|---|---|---|---|
| *Intercept* | 0.142282 | 0.037192 | 3.826 | 1.36 e-04*** |
| *Year_presence* | –0.037273 | 0.005909 | –6.308 | 3.71 e-10*** |

**Table S2.** Summary of the linear model assessing the effect of years since invasion on snake body condition. Body condition was calculated as the residuals of a log–log regression of body mass on snout–vent length (SVL). The model tested whether body condition declines with time since invasion. Model output: Residual standard error = 0.7703 on 1,520 degrees of freedom; $R^2 = 0.0255$; Adjusted $R^2 = 0.0249$; F-statistic = 39.79 on 1 and 1,520 DF; $p < 0.001$.

***(a) Presence-absence of snakes (Zero-inflated model)***

| *Variable* | Odds ratio [95% CI] | $\chi^2$ | p |
|---|---|---|---|
| *(Intercept)* | 0.10 [-0.31, 0.50] | - | 0.64 |
| *Years from invasion* | -1.58 [-2.32, -0.84] | 17.60 (df = 1) | < 0.001 |

***(b) Trap performance (Gamma model)***

| *Variable* | Odds ratio [95% CI] | $\chi^2$ | p |
|---|---|---|---|
| *(Intercept)* | -3.62 [-3.83, -3.41] | - | < 0.001 |
| *Years from invasion* | -0.31 [-0.54, -0.07] | 6.56 (df = 1) | 0.010 |
| ***Random effects*** | | | |
| $\sigma^2$ | 0.33 | ICC | 0.33 |
| | | Marginal $R^2$ | 0.006 |
| $\tau_{00}$ | $0.10_{Site}$ | Conditional $R^2$ | 0.364 |
| | $0.08_{Year}$ | AICc | -11954.20 |

**Table S3.** Results from the zero-inflated Gamma model assessing the effects of year pf invasion on snake presence-absence (a) and trap performance (b). The zero-inflated component (a) models the probability of snake absence, while the Gamma component (b) models trap performance. Predictor variables only include ‘years from invasion’. The table reports odds ratios [95% confidence intervals], Wald chi-square statistics ($\chi^2$) with degrees of freedom, and p-values. Random effects include variance components ($\sigma^2$, $\tau_{00}$) at the ‘site’ and ‘sampling year’ levels, as well as intra-class correlation (ICC), marginal and conditional $R^2$ values, and model selection criterion (AICc).

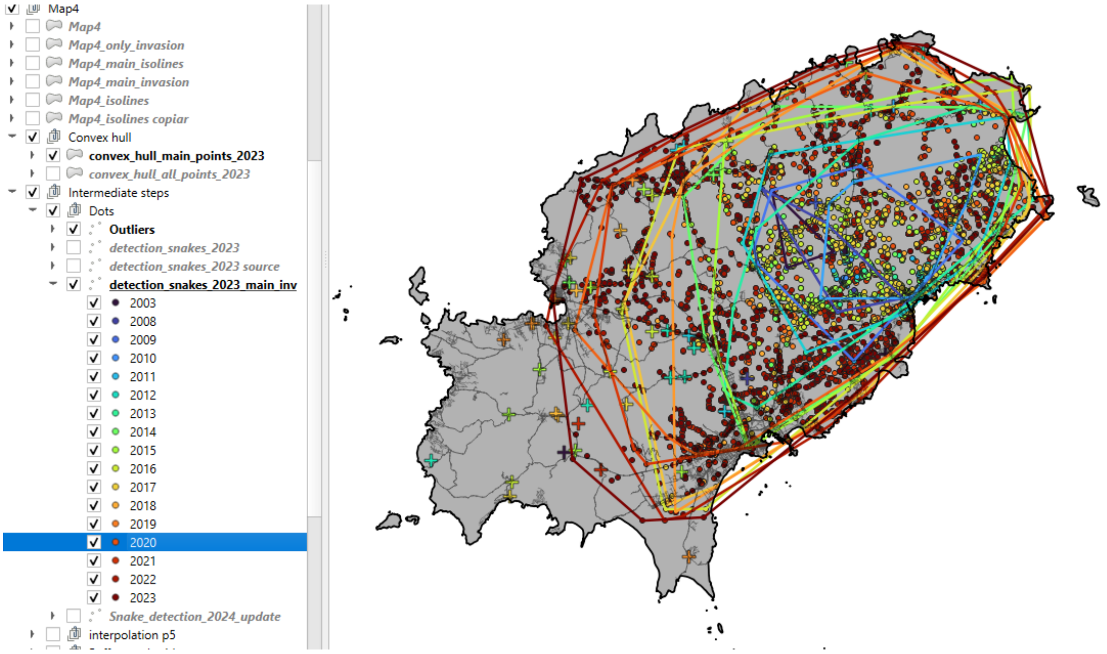


**Figure S1**. Spatiotemporal distribution of snake detections across Ibiza (2003–2023). Each dot represents a georeferenced record of the horseshoe whip snake presence colored by year of first detection. Polygons show yearly convex hulls of invasion extent based on confirmed detections for each year, illustrating the outward progression$f$s2 of the invasion front.

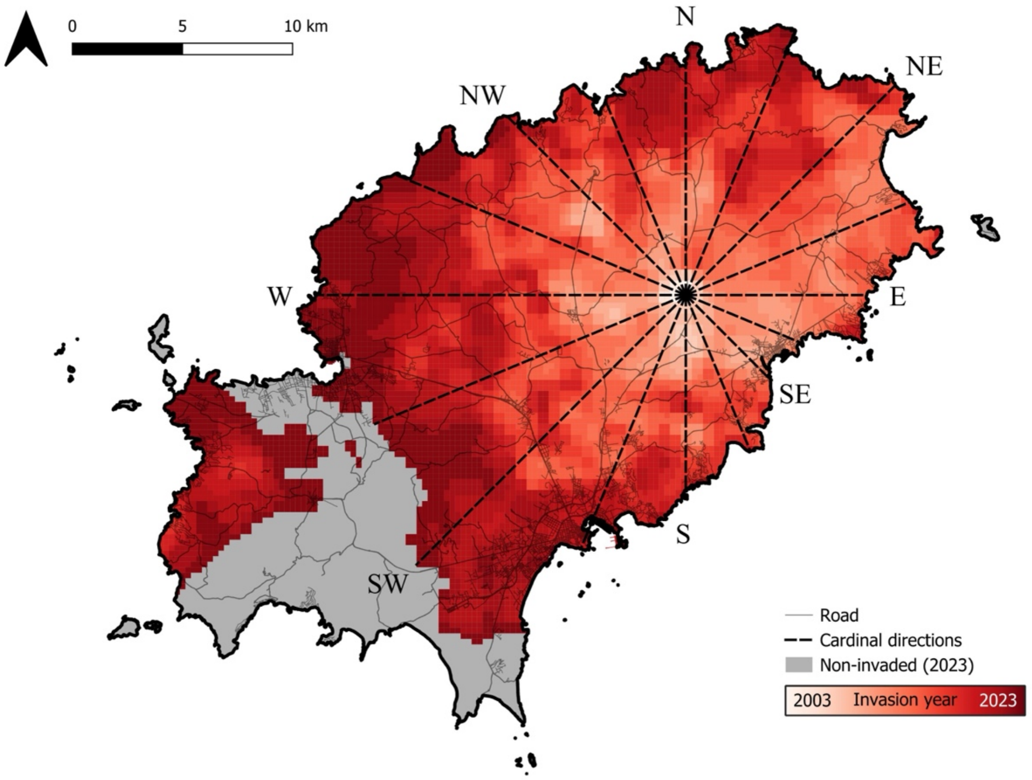


**Figure S2.** Cardinal direction framework used to calculate the directional expansion of the snake invasion across Ibiza.

The map shows the estimated year of *Hemorrhois hippocrepis* presence across the island, based on interpolated invasion data. The 16 black dashed lines represent cardinal and intercardinal axes radiating from the inferred introduction point. Along each axis, we recorded the annual distance reached by the invasion front from 2003 to 2023. These measurements were used to calculate directional expansion rates, assess temporal variation in spread speed, and detect changes in invasion dynamics over time. Grey areas indicate zones without confirmed snake presence as of 2023.

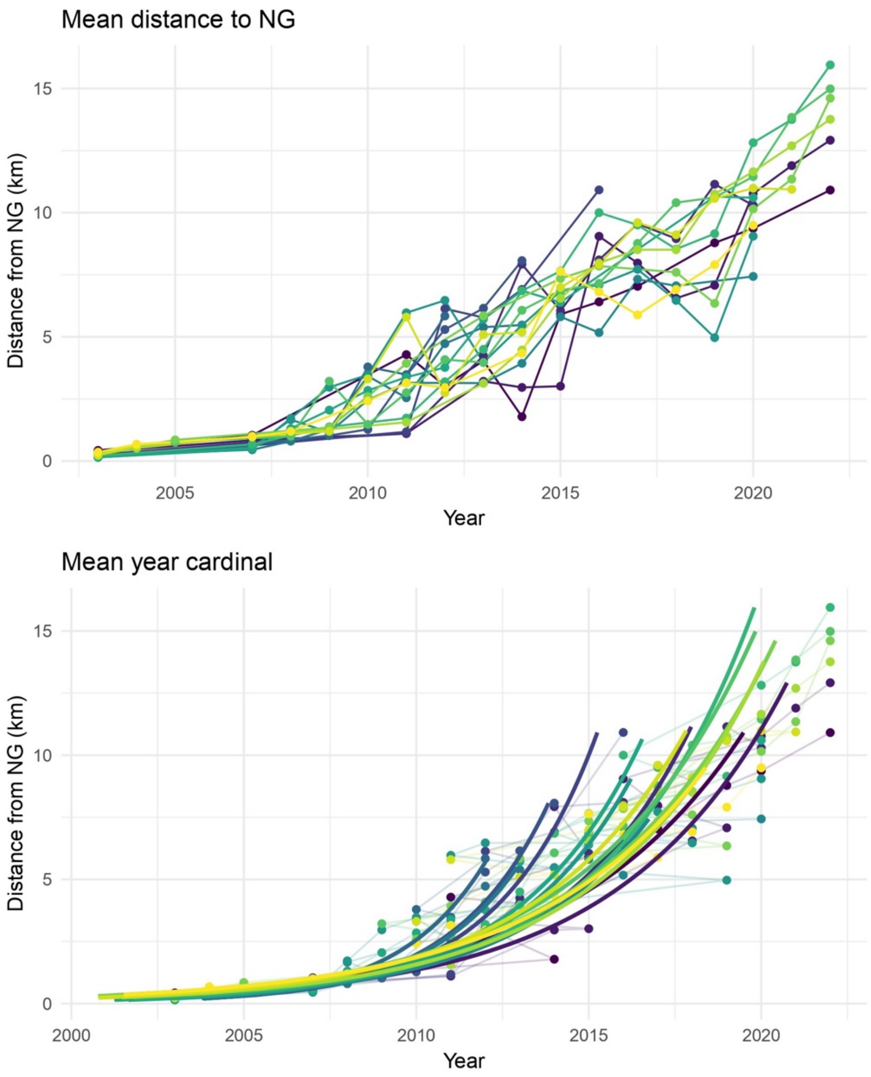


**Figure S3.** Mean invasion-front distance for each cardinal direction, stopping when the coastline is reached (if applicable). The top panel shows points connected according to cardinal direction, while the bottom panel displays the exponential trend for each direction.

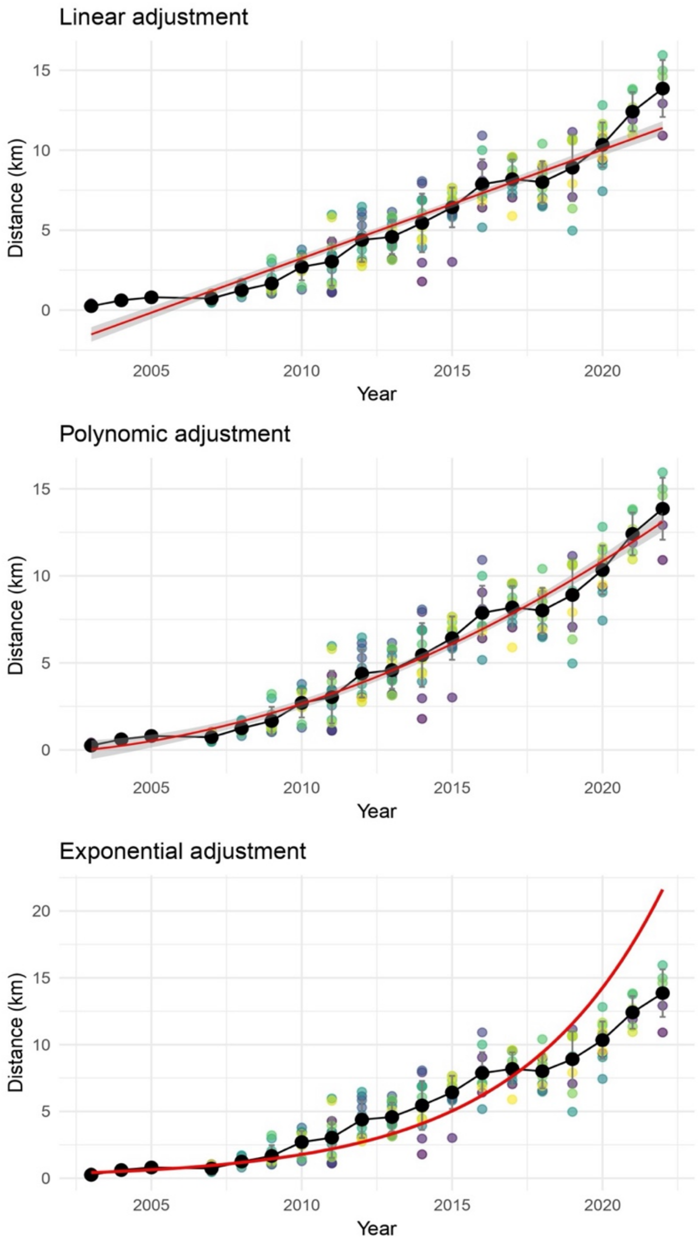


**Figure S4.** Linear, polynomial, and exponential model fits describing the spatial expansion of the invasive horseshoe whip snake across Ibiza from 2003 to 2024. Black dots with error bars represent the annual average distance reached by the invasion front, calculated across 16 radial axes from the introduction point. Colored points show mean raw distance data for each axis and year. The top panel shows the linear fit, the middle panel shows the second-order polynomial fit, and the bottom panel shows the exponential fit, illustrating how non-linear models better capture the accelerating spread of the invasion over time.
